# Regulation of REM Sleep Onset and Homeostasis by Preoptic Glutamatergic Neurons

**DOI:** 10.1101/2024.10.03.616573

**Authors:** Alejandra Mondino, Amir Jadidian, Brandon Toth, Viviane S. Hambrecht-Wiedbusch, Leonor Floran-Garduno, Duan Li, A. Kane York, Pablo Torterolo, Dinesh Pal, Christian Burgess, George A. Mashour, Giancarlo Vanini

## Abstract

The preoptic area of the hypothalamus is key for the control of sleep onset and sleep homeostasis. Although traditionally considered exclusively somnogenic, recent studies identified a group of preoptic glutamatergic neurons that promote wakefulness. Specifically, our previous investigations demonstrated that chemogenetic stimulation of glutamatergic neurons within the medial-lateral preoptic area (MLPO_VGLUT2) promotes wakefulness, fragments non-rapid eye movement sleep (NREMs), and suppresses REM sleep (REMs). This evidence is further supported by recent work showing that preoptic glutamatergic neurons are activated during microarousals that fragment sleep in response to stress, and optogenetic stimulation of these neurons promotes microarousals and wakefulness. Thus, while the wake-promoting function of MLPO_VGLUT2 is clear, their role in sleep homeostasis has not been assessed. We tested the hypothesis that MLPO_VGLUT2 are wake-active, and their activation will increase wakefulness and disrupt sleep homeostasis via projections to arousal-promoting systems. Using fiber photometry, we found that MLPO_VGLUT2 were highly active during REMs, wakefulness and brief arousals, and remained minimally active during NREMs. Chemogenetic stimulation of MLPO_VGLUT2 inhibited REMs onset and suppressed the REMs homeostatic response after total sleep deprivation. Chemogenetic inhibition of MLPO_VGLUT2 increased REMs time (during the light phase only) but did not influence REMs and NREMs homeostasis. Anterograde projection mapping revealed that MLPO_VGLUT2 innervate central regions that promote wakefulness and inhibit REMs. We conclude that MLPO_VGLUT2 powerfully suppress REMs and that exogenous —and possibly pathologic— activation of these neurons disrupts REMs recovery, presumably by directly or indirectly activating REMs-inhibitory mechanisms.

**SIGNIFICANCE STATEMENT:** The preoptic area of the hypothalamus has been extensively studied and its role in sleep regulation is well-established. Importantly, recent work identified a group of preoptic glutamatergic neurons (MLPO_VGLUT2) that are wake-active and promote wakefulness. However, whether these neurons influence sleep homeostasis remains unknown. We demonstrate that MLPO_VGLUT2 are maximally active during REM sleep (REMs), wakefulness and brief arousals from sleep, and innervate wake-promoting and REMs-inhibitory regions. MLPO_VGLUT2 stimulation inhibits REMs and REMs rebound after sleep deprivation, whereas their inactivation increases REMs but does not alter REMs homeostatic response. We thus identified a preoptic mechanism that powerfully suppresses REMs, which we propose may engage during normal sleep-to-wake transitions to block REMs intrusions into subsequent wakefulness.

## INTRODUCTION

Ample evidence from investigations conducted across different species indicates that the preoptic area of the hypothalamus is essential for sleep generation and sleep homeostasis. Experimental lesions (Nauta, 1946; Sallanon et al., 1989; John et al., 1994; Lu et al., 2000; Eikermann et al., 2011) or inhibition (Lin et al., 1989; Alam and Mallick, 1990; Benedetto et al., 2012) of the preoptic area produces insomnia, whereas pharmacologic stimulation increases sleep (Ticho and Radulovacki, 1991; Mendelson, 2000). Selective stimulation of preoptic GABAergic-galaninergic and glutamatergic-nitrergic neurons increases non-rapid eye movement sleep (NREMs) (Chung et al., 2017; Harding et al., 2018; Kroeger et al., 2018; Vanini et al., 2020; Mondino et al., 2021; Machado et al., 2022). Within the preoptic area, a subset of neurons increases their activity during wakefulness with increased sleep pressure (Gvilia et al., 2006; Alam et al., 2014; Gvilia et al., 2017). Ablation of galaninergic neurons in the mouse lateral preoptic area reduces electroencephalographic delta power rebound during recovery sleep, suggesting an important role for these cells in sleep homeostasis (Ma et al., 2019). Additionally, preoptic GABAergic neurons that project to the tuberomammillary nucleus are mostly active during periods of increased rapid eye movement sleep (REMs) pressure, and their inhibition reduces REMs time and blocks the homeostatic rebound after REMs restriction (Maurer et al., 2024). These data are consistent with previous reports suggesting a specific role for preoptic neurons in REMs control (Lu et al., 2002).

Although traditionally considered exclusively somnogenic, evidence from several independent studies indicates that neuronal subpopulations in the preoptic area also promote wakefulness. Single-unit (Koyama and Hayaishi, 1994; Takahashi et al., 2009) and fiber photometry (Smith et al., 2024) recordings revealed that a group of preoptic neurons is active during brief arousals and wake states. Stimulation of subsets of preoptic glutamatergic (Vanini et al., 2020; Mondino et al., 2021; Smith et al., 2024) and tachykinin-1 (Reitz et al., 2020) neurons promotes brief arousals and increases wakefulness. We recently demonstrated that chemogenetic stimulation of glutamatergic neurons within the ventral portion of the medial and lateral preoptic area (MLPO_VGLUT2) produces (1) brief electroencephalographic and behavioral arousals from sleep that increase the total time spent in wakefulness, (2) state instability that causes a robust fragmentation of NREMs, (3) REMs suppression, (4) altered cortical oscillation and dynamic patterns consistent with a lighter NREMs state, and (5) body cooling (Mondino et al., 2021). Collectively, these data demonstrate that a group of preoptic glutamatergic neurons promotes wakefulness, fragments NREMs, and suppresses REMs.

Unlike studies of preoptic GABAergic and galaninergic neurons, the role of preoptic wake-promoting glutamatergic neurons in sleep homeostasis has not been assessed. Furthermore, the downstream regions by which these preoptic glutamatergic neurons achieve their arousal-promoting effects remain to be identified. This study tested the hypothesis that MLPO_VGLUT2 are wake-active, and their selective activation will disrupt sleep homeostasis and increase wakefulness via direct projections to arousal-promoting systems. Using fiber photometry, we found that the activity of MLPO_VGLUT2 during spontaneous sleep-wake states was highest during REMs and wakefulness, and lowest during NREMs. Chemogenetic activation of these preoptic neurons inhibited REMs onset and suppressed the REMs homeostatic response to total sleep deprivation. Chemogenetic inhibition of MLPO_VGLUT2 increased the time spent in REMs (during the light phase only) but did not alter REMs recovery after sleep deprivation. Anterograde tracing studies revealed that MLPO_VGLUT2 send efferent projections to central regions that promote wakefulness and inhibit REMs, suggesting complex circuitry that modulates sleep-wake behavior rather than mere direct influences on cortical arousal. Thus, our study revealed that MLPO_VGLUT2 powerfully suppress REMs occurrence and that exogenous —and possibly pathologic— activation of these neurons disrupts REMs recovery, presumably through projections to arousal-promoting and REMs-inhibitory regions.

## MATERIALS AND METHODS

### Mice

We studied transgenic Vglut2-IRES-Cre (Slc17a6^tm2(cre)Lowl^/J) mice. Breeding pairs were purchased from The Jackson Laboratory (stock #016963) and bred at the University of Michigan animal care facility to establish and maintain the colony. A group of Vglut2-IRES-Cre mice was crossed with eGFP-L10a reporter mice that express a green fluorescent protein in glutamatergic neurons. These Vglut2-Cre;eGFP-L10a mice (Vong et al., 2011; Beekly et al., 2020) were used as controls in photometry experiments. All mice were genotyped (Transnetyx) before weaning. Adult male and female mice (n = 35; 16 to 20 weeks old, 16 to 24 g at the time of surgery) were housed in groups under a 12-h light/dark cycle (lights on at 6:00 AM) at constant temperature and humidity (22⁰ C and 50-55%) levels. Food and water were supplied ad libitum.

### Viral Vectors and Designer Receptor Agonist

To study the activity of preoptic glutamatergic (Vglut2+) neurons during spontaneous sleep-wake states, we used the adeno-associated viral vector AAV8-Syn-FLEX-jGCaMP7f-WPRE (Dr. Douglas Kim; Addgene, Cat.# 104492-AAV8) for Cre-recombinase-mediated expression of the calcium sensor protein GCaMP7f (Dana et al., 2019). For chemogenetic stimulation and inhibition of preoptic glutamatergic neurons, we used the Cre-recombinase-dependent adeno-associated viral vectors AAV5-hSyn-DIO-hM3D(Gq)-mCherry and AAV8-hSyn-DIO-hM4D(Gi)-mCherry (Dr. Bryan Roth; Addgene, Cat.# 50459-AAV5 and Cat.# 44362-AAV8) (Krashes et al., 2011). We performed conditional anterograde tracing to investigate the projections of glutamatergic neurons in the MLPO region. The AAV8-EF1a-double floxed-hChR2(H134R)-mCherry-WPRE-HGHpA (Dr. Karl Deisseroth; Addgene, Cat.# 20297-AAV8) was bilaterally injected into the MLPO of Vglut2-cre mice for Cre-inducible expression of channelrhodopsin-2 conjugated with mCherry. These viral vectors had a titer concentration of 7.8 X 10^12^, 1.9 X 10^13^, and 1.9 X 10^13^ genome copies per ml, respectively.

The designer receptor agonist Clozapine-N-oxide (CNO; Cat.# C0832-5MG) and the solvent dimethyl sulfoxide (DMSO) were purchased from Sigma-Aldrich (St. Louis, MO, USA). CNO was dissolved in sterile saline containing DMSO. A stock solution of CNO (0.1 mg/ml in sterile saline and 0.5% DMSO) was made, divided in aliquots, and stored at −20°C for subsequent use. For each experiment, one aliquot of the CNO stock solution was allowed to thaw at room temperature and was protected from light until use. The injection volume was 0.1 ml per 10 g of mouse weight.

### Surgical Procedures for Vector Injection, Fiber Photometry Recordings, and Sleep Studies

As described previously (Vanini et al., 2020; Mondino et al., 2021), mice were anesthetized in an induction chamber with 5.0% isoflurane (Hospira, Inc., Lake Forest, IL, USA) in 100% O2 and then injected with a non-steroidal anti-inflammatory drug (5 mg/kg carprofen or 2 mg/kg meloxicam, subcutaneous) for preemptive analgesia. Thereafter, mice were positioned within a Kopf Model 962 stereotaxic frame fitted with a mouse adaptor (Model 922) and a mouse anesthesia mask (Model 907) (David Kopf Instruments, Tujunga, CA, USA). The concentration of isoflurane (monitored by spectrometry using a Cardiocap™/5; Datex-Ohmeda, Louisville, CO, USA) was then reduced to 1.6-2.0% for the remainder of the surgical procedure. Core body temperature was maintained at 37-38° C throughout surgery using a water-filled pad connected to a heat pump (Gaymar Industries, Orchard Park, NY, USA). Under sterile conditions, mice received a microinjection of AAV8-Syn-FLEX-jGCaMP7f-WPRE (36 nl), or bilateral microinjections of AAV5-hSyn-DIO-hM3D(Gq)-mCherry (36 nl), AAV8-hSyn-DIO-hM4D(Gi)-mCherry (36 nl), or AAV8-EF1a-double floxed-hChR2(H134R)-mCherry-WPRE-HGHpA (36 nl) into the MLPO region using the following stereotaxic coordinates (Paxinos and Franklin, 2001): 0.10 mm anterior to bregma, ±0.5 mm lateral to the midline, and 5 mm below the dura. The vector injection (5 nl/min) was performed using a Hamilton Neuros Syringe 7000 (5 μL; Hamilton Company, Reno, NV, USA) mounted on a microinjection syringe pump connected to a digital Micro2T controller (Model UMP3T-2; World Precision Instruments, Sarasota, FL, USA). After each injection, the syringe was kept in position for at least 5 minutes to avoid vector reflux. The delivery of isoflurane was discontinued, and then mice were placed under a heating lamp and monitored until fully ambulatory. Analgesia was maintained for a minimum of 48 hours post-surgery. Mice were allowed a minimum of 5 days to recover from surgery.

Two to three weeks after the vector injection, mice were implanted with stainless-steel screw electrodes (8IE3632016XE, E363/20/1.6/SPC; Plastics One, Roanoke, VA, USA) to record the electroencephalogram (EEG) from frontal (1.5 mm anterior to bregma and ±2.0 mm relative to the midline), and occipital (3.2 mm posterior to bregma and ±3.0 mm relative to the midline) regions. A screw electrode implanted over the cerebellum served as reference for monopolar EEG recordings, and two wire electrodes (8IE36376XXXE, E363/76/SPC; Plastics One) implanted bilaterally in the neck muscles were used to record the electromyogram (EMG). All electrodes were then inserted into a six-pin electrode pedestal (MS363; Plastics One).

During the same surgery, mice to be studied with photometry were also implanted with an optical fiber (Product # CFML15L10; 400-μm diameter core, OD 1.25 mm, NA 0.50, length 10 mm; ThorLabs) over the MLPO region at coordinates: 0.10 mm anterior to bregma, ±0.6 mm lateral to the midline, and 5.5 mm below the dura. The pedestal, recording electrodes, and the optical fiber (when implanted) were cemented to the skull using dental acrylic (Fast Cure Powder/Liquid, Product# 335201; GC America, Inc., Alsip, IL, USA). Postoperative analgesia and care were conducted as described above for vector injection.

### Experimental Design of Sleep Experiments

All experiments and procedures were approved by the Institutional Animal Care and Use Committee and the Institutional Biosafety Committee, and were conducted in accordance with the Guide for the Care and Use of Laboratory Animals (Ed 8, National Academies Press, Washington, DC, 2011). After recovering from surgery, mice were habituated to handling, intraperitoneal injections, tethering to the EEG-EMG recording cables, and sleep recording chambers (Raturn BASi System; West Lafayette, IN, USA) for at least 5 days before experiments.

### In vivo fiber photometry recordings during spontaneous sleep-wake states

Two to three weeks after surgery, mice were connected to an EEG/EMG recording tether and a fiber optic patch cable, and given 5-7 days to habituate to the recording environment. Mice were considered fully habituated after having built a nest and were comfortably feeding and drinking in the recording chamber. Mice were able to move freely within the recording chamber while outfitted with both optic fibers and EEG/EMG tethers. Fiber optic patch cables (0.8-1 m long, 400 μm diameter; Doric Lenses) were firmly attached to the implanted fiber optic cannulae with zirconia sleeves (Doric Lenses). LEDs (Plexon; 473 nm) were set such that a light intensity of <0.1 mW entered the brain; light intensity was kept constant across sessions for each mouse. Emission light was passed through a filter cube (Doric) before being focused onto a sensitive photodetector (2151, Newport). Signals were digitized at 60 Hz using PyPhotometry (Akam and Walton, 2019), which allows for pulsed delivery of light, minimizing bleaching throughout 6-hour recordings. For time normalization of photometry signals across bouts of a given state, photometry signals were first downsampled to 10 Hz, then linearly interpolated to the same length.

EEG and EMG signals recorded during photometry experiments were digitized at 1000 Hz, with a 0.3-100 Hz bandpass filter applied to the EEG and a 30-100 Hz bandpass filter applied to the EMG (ProAmp-8, CWE Inc.), with a National Instruments data acquisition card and collected using a custom MATLAB script. EEG/EMG signals were notch-filtered at 60 Hz to account for electrical interference from the recording tether. Prior to analysis, EEG signals were high-pass filtered at 1 Hz to remove low-frequency artifacts from the recordings. Recordings were 12 hours long and contained the second half of the dark cycle (ZT18 – ZT24) and the first half of the light cycle (ZT0 – ZT06). Polysomnographic signals were analyzed using AccuSleep, an open-source sleep scoring algorithm in MATLAB, and verified by manual inspection. Behavioral states were scored in 5-s epochs as either wake, NREMs, or REMs (a detailed description of scoring criteria is provided below in “*Sleep Data Acquisition and Analysis*” section). Brief arousals were defined as transitions from NREMs to wake that lasted <20 s (Franken et al., 1999; Cardis et al., 2021; Park et al., 2021). In subsequent photometry analysis, for primary state transitions, only transitions in which at least 20 s of a state occurred on either side of the transition were included.

### Chemogenetic stimulation and inhibition of MLPO glutamatergic neurons

Each set of chemogenetic studies used a within-subject design so that each mouse underwent all treatments and served as its own control. Mice received randomized intraperitoneal injections of vehicle or CNO (1 mg/kg), counterbalanced across cohorts, and were allowed a washout period of at least 3 days after chemogenetic experiments. Baseline EEG and EMG recordings used to assess the effects of sleep deprivation on sleep-wake states were obtained for 6 hours beginning at ZT06 (1 PM). On a separate day, mice were maintained awake for 6 hours beginning at the onset of the light phase (ZT0 = 7 AM) by providing novel objects, nesting materials, tapping on the side of the cage, or by gentle stimulation of the tail using a pencil-sized paintbrush whenever the mouse appeared immobile and drowsy, or a sleep attempt was observed (Vanini, 2016; Hambrecht-Wiedbusch et al., 2017). Thirty minutes prior to the end of sleep deprivation, mice received an injection of vehicle or CNO. After sleep deprivation, the EEG and EMG were recorded for 6 hours (ZT06 to ZT12). In a separate set of experiments, mice received an injection of vehicle or CNO (6:30 PM; 30 minutes before the onset of the dark phase), followed by EEG and EMG recordings for 6 hours (ZT12 to ZT18). These two sets of experiments were designed to assess the role of MLPO_VGLUT2 in the regulation of sleep-wake states in response to a higher (post-sleep deprivation) or lower (night) homeostatic sleep drive, respectively. Next, chemogenetic inhibition experiments were conducted during the light phase (period of highest sleep drive), the dark phase (period of lowest sleep drive), and after sleep deprivation, to determine whether spontaneous activation of MLPO_VGLUT2 causally contributes to regulating sleep-wake states and sleep homeostasis. Thus, experiments in mice expressing inhibitory designer receptors in MLPO_VGLUT2 followed the same protocols described above for stimulation experiments.

### Sleep Data Acquisition and Analysis

EEG and EMG signals were amplified (×1000) and digitized (sampling rate = 1024 Hz), respectively, with a model 1700 AC amplifier (A-M Systems) and a Micro3 1401 acquisition unit and Spike2 software (Cambridge Electronic Design); a 60 Hz filter was applied to the recordings when electrical noise was evident. EEG signals were bandpass filtered between 0.1 and 500 Hz and EMG signals were bandpass filtered between 10 and 500 Hz. Mouse behaviors were video recorded in synchrony with the electrophysiologic recordings in Spike2. States of wakefulness, NREMs, and REMs were manually scored in 5 s epochs using Spike2 software. Scoring results were verified by an independent investigator who was blinded to the treatment condition.

Wakefulness was defined by low-amplitude, high-frequency EEG activity accompanied by an active EMG characterized by high tone with phasic movements. NREMs was defined by high-amplitude, low-frequency EEG waveforms along with low muscle tone. REMs was defined by low-amplitude, high-frequency EEG signals with a prominent theta rhythm and muscle atonia reflected in the EMG. Total time spent in wakefulness, NREMs, and REMs, the average duration and number of bouts for each vigilance state, were analyzed over the 6 h recording period, as well as in 1 h bins to assess the mean duration of the treatment effect. The mean latency to NREMs and REMs onset was compared between treatment conditions.

### Power Spectra and Sleep Spindle Detection Analysis

After chemogenetic stimulation experiments, the EEG spectral power was computed using the multitaper method in Chronux analysis toolbox (Mitra and Bokil, 2008) with a window length = 5 s without overlap, time-bandwidth product = 2, and number of tapers = 3. For each mouse in each treatment group, the mean power spectrum during wakefulness, NREMs, and REMs was computed by averaging the power spectra across all available windows in the frontal channels. Epochs containing artifacts and state transitions were not included in the analysis. Power values were calculated for slow oscillations (0.5-1 Hz), delta (1-4 Hz), theta (4-9 Hz), sigma (9-15 Hz), beta (15-30 Hz), low gamma (30-45 Hz), and high gamma (75-115 Hz) frequency bands.

Additionally, power values were normalized for each mouse as the mean power of each frequency band divided by the sum of the power in all bands (i.e., total power). The number and duration of sleep spindles was calculated using a previously validated MATLAB script, kindly provided by Dr. James McNally (VA Boston Healthcare System and Harvard Medical School) (Uygun et al., 2019).

### Anterograde tracing studies

For a qualitative assessment of projection targets of MLPO_VGLUT2, Vglut2-Cre mice (n = 5) received bilateral injections of a vector for Cre-inducible expression of channelrhodopsin conjugated to the mCherry reporter. Six to eight weeks after vector injection, mice were transcardially perfused and the brains were removed, sectioned coronally, and immunostained for mCherry (perfusion and immunohistochemistry protocols are described below). Only mice with confirmed expression of channelrhodopsin in the MLPO region were included in the analysis. For mapping the projections of MLPO_VGLUT2, rostrocaudal series of brain sections (anterior-posterior coordinates +3.0 to -7.5 mm relative to bregma (Paxinos and Franklin, 2001) were systematically examined at 20X and 40X magnification using an Olympus BX43 fluorescence microscope (Olympus America). The brain areas containing labeled axons were identified by matching the brain section with the corresponding coronal section in a mouse brain atlas (Paxinos & Franklin, 2001).

### Immunohistochemistry and Histology

After completing data collection, mice were deeply anesthetized and transcardially perfused with 0.1 M phosphate buffered saline (PBS), pH 7.4, followed by 5% formalin for 10 min using a MasterFlex perfusion pump (Cole Palmer). The brains were then extracted and postfixed in 5% formalin overnight at 4°C. Subsequently, brains were cryoprotected with 20% sucrose in PBS for 1-2 days, and then transferred to 30% sucrose for an additional 2 days. Thereafter, brains were frozen in Tissue–Plus (Fisher Healthcare) and sectioned coronally at 40 µm using a cryostat (CM1950, Leica Microsystems). Brain sections that contained the MLPO region (or rostrocaudal series of sections for the projection mapping study) were collected and blocked in PBS containing 0.25% Triton X-100 and 3% normal donkey serum (Vector Laboratories) for 60 min at room temperature. The tissues were then incubated overnight at room temperature in primary antiserum (rat monoclonal anti-mCherry 1:50,000; Thermo Fisher Scientific, catalog #<u>M11217</u>), followed by incubation for 2 h in a donkey anti-rat secondary antiserum (1:750; AlexaFluor-594, Thermo Fisher Scientific, catalog #A-21209). Last, brain sections were mounted on glass slides, and coverslipped with SlowFade Diamond (<u>S36972</u> or <u>S36973</u>;Invitrogen). The correct localization of the designer receptors was confirmed using fluorescence microscopy (BX43, Olympus America), with the aid of a mouse brain atlas (Paxinos and Franklin, 2001).

### Statistical Analysis

Data analysis was performed with PRISM version 9.3 (GraphPad Software), with input from the Consulting for Statistics, Computing & Analytics Research unit, University of Michigan. All data were tested for normality and are presented as mean ± standard error of the mean (SEM); a p < 0.05 was considered statistically significant. Analysis of the activity of MLPO_VGLUT2 using fiber photometry was conducted using a one-way analysis of variance (ANOVA), followed by Tukey’s multiple comparisons test. Analysis of sleep-wake parameters and EEG power was conducted using a two-way, repeated-measures ANOVA or a mixed-effects model (in cases of missing data), both with a Geisser-Greenhouse correction followed by Šídák’s multiple comparisons test. Statistical comparison of EEG power in selected frequencies between baseline and sleep deprivation, as well as sleep latencies was analyzed with a paired t-test or the non-parametric Wilcoxon test. Specific statistical analyses used in each study are provided in the respective description of the results.

## RESULTS

### Medial-Lateral Preoptic Glutamatergic Neurons Are Most Active During REM sleep

In our previous study, we showed that chemogenetic activation of MLPO_VGLUT2 promoted arousal (Vanini et al., 2020; Mondino et al., 2021). However, the activity pattern of MLPO_VGLUT2 during naturally occurring sleep-wake states has not been investigated. To fill this gap in knowledge, we expressed the Cre-inducible AAV8-Flex-jGCaMP7f in the MLPO region of Vglut2-Cre mice (n = 5) and conducted fiber photometry recordings of these neurons during objectively identified states of sleep and wakefulness (Figure 1A-B). There was a significant state effect in GCaMP-expressing mice (F(2, 8) = 18.79, p = 0.0009), but not in control Vglut2-Cre;eGFP mice (n = 3) that did not receive AAV injection. On average, activity in GCaMP-expressing mice was highest during REMs, mildly elevated during wakefulness, and lowest during NREMs (Tukey’s multiple comparisons test; p = 0.0157 (Wake vs. NREMs) and p = 0.0008 (NREMs vs. REMs), p = 0.0939 (Wake vs. REMs); Figure 1C-D). Within individual bouts of each arousal state, we observed that while the activity in REMs remains consistently elevated, the activity during wakefulness significantly decreases over time (Paired t-test, p = 0.006; Figure 1C-D). Additionally, there was a steep increase in the activity of MLPO_VGLUT2 immediately before the transition from NREMs into REMs and wakefulness. However, while the activity of these neurons plateaued during REMs bouts, it gradually declined during wakefulness bouts (Figure 1E). Consistent with the arousal-promoting effect of these preoptic neurons, transient increases in activity were observed during brief arousals from sleep defined as a period of wakefulness of < 20 s in duration between consecutive NREMs bouts. Importantly, none of the dynamics described here were observed in the preoptic area of control mice expressing native eGFP (Figure 1E).

**Figure 1.**
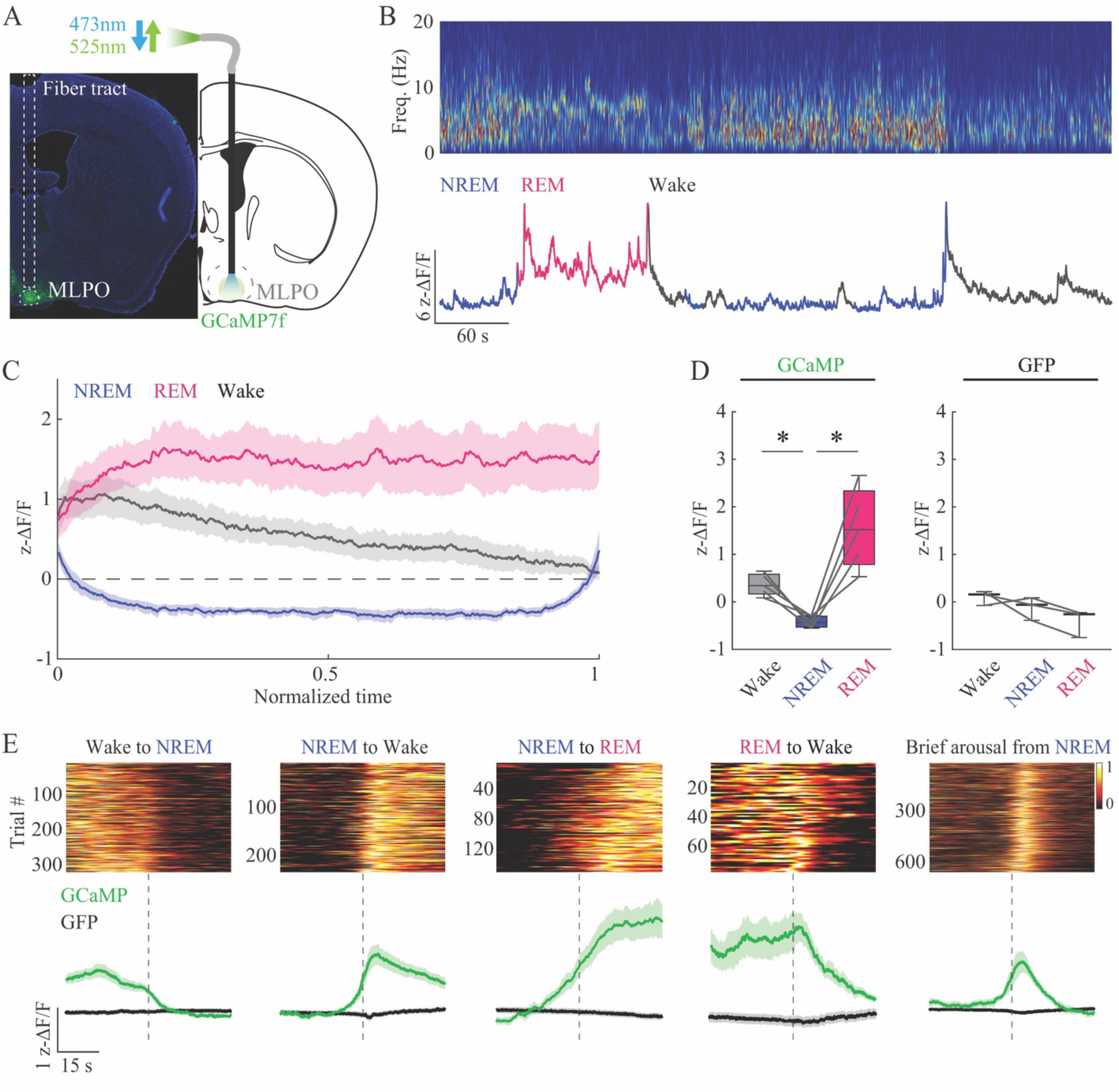
Medial-lateral preoptic glutamatergic neurons are highly active during REM sleep and wakefulness. **A,** Low-magnification photograph shows the expression of the calcium sensor GCaMP7f (green: GCaMP, blue: DAPI staining) in the MLPO of a Vglut2-Cre mouse and a schematic of photometry recording from neurons in the MLPO region of the hypothalamus. **B,** Representative power spectrogram (top) and ΔF/F trace (bottom) across sleep-wake states. **C,** Temporal dynamics of GCaMP fluorescence across bouts of wake, NREM, and REM sleep in normalized time. **D,** Mean fluorescence during wakefulness, NREM, and REM sleep in Vglut2-Cre mice expressing GCaMP7 and control mice expressing GFP in MLPO glutamatergic neurons. **E,** Single-trial transition-onset-triggered time courses of GCaMP fluorescence, sorted by response amplitude at the onset of arousal state transitions (top). Average GCaMP (solid green trace) and GFP (solid black line) fluorescence aligned to arousal state transitions (dashed grey line; bottom). Data are shown as mean ± SEM. *, p < 0.05.

### Chemogenetic Stimulation of Medial-Lateral Preoptic Glutamatergic Neurons Inhibited REM Sleep Homeostasis

Previously published work demonstrated that activation of MLPO_VGLUT2 – during the light phase – increased wakefulness, fragmented NREMs, and prevented REMs generation (Vanini et al., 2020; Mondino et al., 2021). The current study aimed to investigate the effects of the activation of these neurons on the regulation of sleep-wake states under high or low homeostatic sleep drive conditions. Thus, recordings of sleep-wake states were obtained from n = 11 mice after sleep deprivation during the light phase and at the onset of the dark phase. As expected, sleep deprivation significantly reduced the time in wakefulness compared to baseline (hours 1, 3, and 6 after sleep deprivation; p value < 0.0001, < 0.0001 and < 0.0001, respectively) and increased the time in NREMs during recovery (hours 1, 2, 3 and 6 after sleep deprivation; p value < 0.0001, 0.0440, < 0.0001 and < 0.0001, respectively). The time in REMs was significantly increased during hours 1, 3, and 6 after sleep deprivation (p = 0.0002, < 0.0001 and < 0.0001); there was a reduction in REMs time during hour 2 (p = 0.0111). Additionally, there was a significant increase in the mean frontal power (average of hours 0 to 2 after sleep deprivation from n = 10 mice) in slow oscillations (t = 4.988; df = 9; p = 0.0008) and delta (t = 2.258; df = 9; p = 0.0252) frequencies during NREMs, reduced NREM theta power (t = 3.582; df = 9; p = 0.0059), and increased power of high-gamma during wakefulness (t = 2.265; df = 9; p = 0.0497); only significant changes for NREMs and wakefulness EEG power are reported. There was no significant change in frontal EEG power during REMs rebound after sleep deprivation.

We previously validated the designer receptor hM3Dq by demonstrating the activation (i.e., cFos expression) of MLPO_VGLUT2, and that the dose of CNO used in the present study did not alter sleep-wake states in control mice without hM3Dq expression (Vanini et al., 2020; Mondino et al., 2021). To determine whether the activation of MLPO_VGLUT2 influences sleep homeostasis, n = 11 mice were sleep deprived, and then received an injection of vehicle or CNO prior to the sleep-wake recording. The statistical analysis of these data was conducted using a two-way, repeated-measures analysis of variance (ANOVA) with a Geisser-Greenhouse correction followed by Šídák’s multiple comparisons test. Figure 2 illustrates the effects of stimulation of MLPO_VGLUT2 on sleep-wake parameters after sleep deprivation. Figure 2A shows representative examples of EEG and EMG recordings from a Vglut2-Cre mouse during states of REMs, NREMs, and wakefulness. Histological analysis confirmed that the expression of the designer excitatory receptors HM3Dq in glutamatergic neurons was localized to the medial-lateral preoptic region of the hypothalamus (Figure 2B). There was a significant time effect, treatment condition (CNO), and treatment × time interaction for REMs (F(5,120) = 7.024; p < 0.0001, F(5,120) = 1450; p < 0.0001 and F(5,120) = 5.08; p = 0.0003), NREMs (F(5,120) = 22.80; p < 0.0001, F(5,120) = 126.9; p < 0.0001 and F(5,120) = 14.55; p < 0.0001), and wakefulness (F(5,120) = 19.62; p < 0.0001, F(5,120) = 7.02; p = 0.0091 and F(5,120) = 15.04; p < 0.0001).

**Figure 2.**
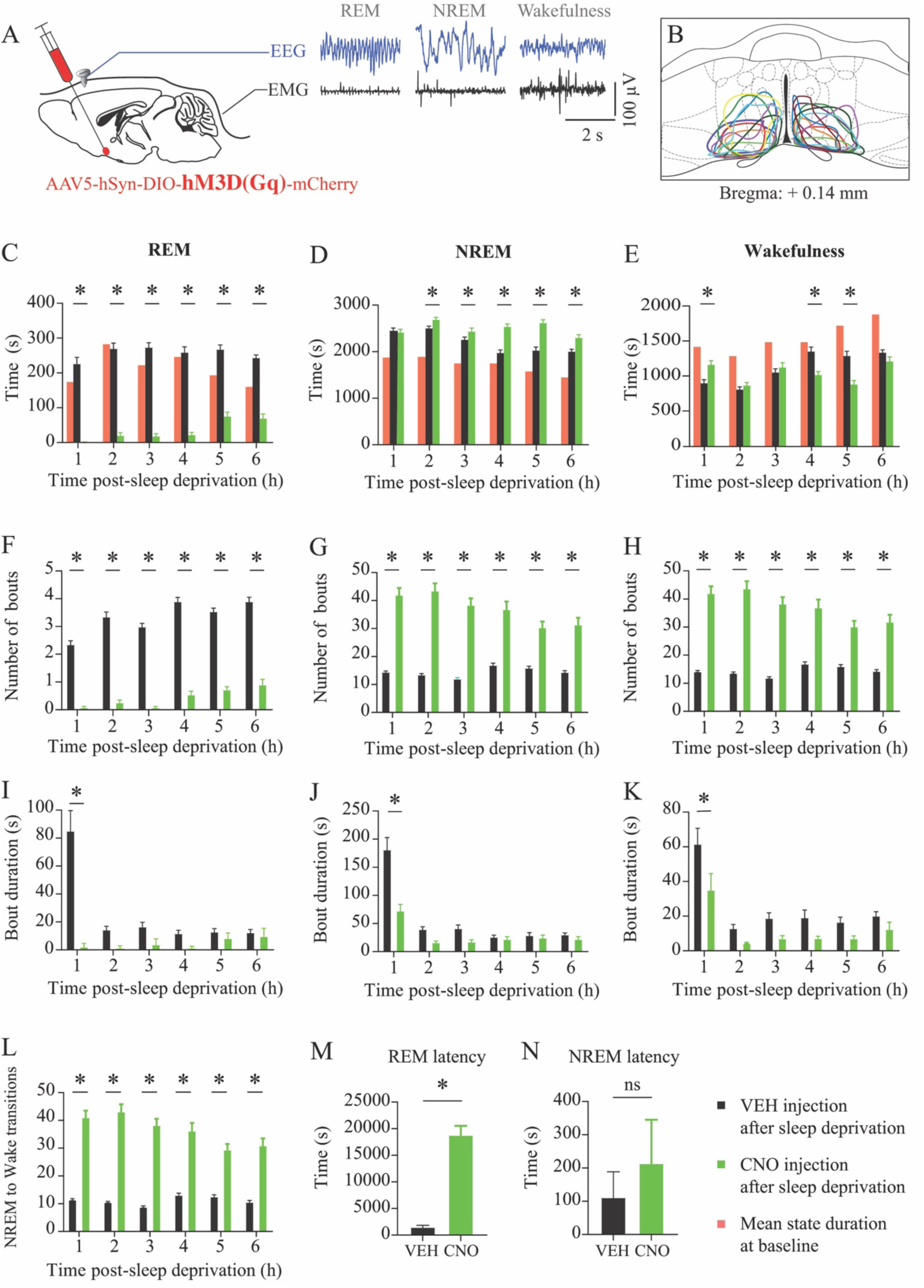
Chemogenetic activation of glutamatergic neurons in the medial-lateral preoptic region inhibited REM sleep homeostatic recovery after sleep deprivation. **A,** Schematic depiction of the injection of a Cre-dependent adeno-associated virus for expression of the excitatory designer receptor hM3Dq into the medial-lateral preoptic region of Vglut2-Cre mice. Representative recordings of the electroencephalogram (EEG) and electromyogram (EMG) obtained from the same Vglut2-Cre mouse during rapid eye movement (REM) sleep, non-rapid eye movement (NREM) sleep, and wakefulness. **B,** Depiction of the vector injection area, represented on a coronal schematic of the preoptic area modified from a mouse brain atlas (Paxinos and Franklin, 2001). **C-E,** Time spent in REM sleep, NREM sleep, and wakefulness, after administration of vehicle (VEH) and clozapine-N-oxide (CNO; 1.0 mg/kg) to sleep deprived mice. For reference, the mean time spent in each state during baseline recordings are indicated by light-red bars. **F-H,** Number of bouts per state. **I-K,** Mean duration of REM sleep, NREM sleep, and wake bouts. **L,** Number of transitions from NREM sleep to wakefulness. **M-N,** Latency to the first REM and NREM sleep bout after CNO administration. Data are shown as mean ± SEM. *, p < 0.05. Asterisks indicate statistical comparisons between vehicle (black bars) and CNO (green bars).

Relative to vehicle, CNO administration caused a significant reduction in the time spent in REMs between hours 1 and 6 after sleep deprivation (p < 0.0001 for each hour bin) (Fig. 2C). CNO administration also significantly increased NREMs duration during hours 2 (p = 0.0269), 3 (p = 0.0405), 4 (p < 0.0001), 5 (p < 0.0001) and 6 (p < 0.0001) after sleep deprivation (the p value for hour 1 was 0.9876) (Fig. 2D). Activation of MLPO_VGLUT2 caused a significant increase in the time in wakefulness during the first hour after sleep deprivation (p = 0.0005) and a reduction of wakefulness duration during hours 4 (p < 0.0001) and 5 (p < 0.0001) after sleep deprivation (Fig. 2E). For reference, the mean hourly time of sleep-wake states during baseline condition (no injection or manipulation) are depicted by the red bars in Figures 2C-D. ANOVA revealed a significant effect on the number of REM (F(1,120) = 1748; p < 0.0001, F(5,120) = 27.97; p < 0.0001 and F(5,120) = 4.550; p = 0.0008), NREM (F(1,120) = 502.1; p < 0.0001, F(5,120) = 3.984; p = 0.0022 and F(5,120) = 6.530; p < 0.0001) and wake (drug, time and interaction: F(1,120) = 504.1; p < 0.0001, F(5,120) = 3.898; p = 0.0026 and F(5,120) = 6.655; p < 0.0001) bouts after sleep deprivation. CNO caused a significant decrease in the number of REMs bouts, and an increase in the number of NREMs and wakefulness bouts after sleep deprivation (p < 0.0001 between hours 1 and 6 for REM, NREM, and wake states) (Figs. 2F-H). Additionally, there was a significant treatment effect (CNO) on bout duration for REMs (F(1,120) = 52.24; p < 0.0001), NREMs (F(1,120) = 41.41; p < 0.0001), and wakefulness (F(1,120) = 25.21; p < 0.0001). CNO injection decreased bout duration during hour 1 post-sleep deprivation in REMs (p < 0.0001), NREMs (p < 0.0001), and wakefulness (p = 0.0002) (Figs. 2I-K). Figure 2L plots the hourly number of NREM to wake transitions during 6 hours after sleep deprivation. ANOVA demonstrated a significant effect of treatment (F(1,120) = 619.4; p < 0.0001) and time (F(5,120) = 4.228; p = 0.0014) as well as a significant treatment x time interaction (F(5,120) = 6.151; p < 0.0001). Post hoc test showed that the number of NREM-to-wake transitions was significantly increased by CNO between hours 1 and 6 post-sleep deprivation (p < 0.0001 in each time bin). CNO injection significantly increased the latency to REMs (mean ± SEM = 1528 ± 322 vs 18828 ± 1713; Wilcoxon, p = 0.0005), whereas the mean latency to NREMs was twice as long after CNO administration but this difference did not achieve statistical significance (111.8 ± 76.82 vs 214.1 ± 130.8; Wilcoxon, p = 0.0527) (Figs. 2M and 2N).

### Activation of MLPO Glutamatergic Neurons Altered the EEG Pattern During Sleep-Wake States in Response to Sleep Deprivation

We recently reported that chemogenetic activation of MLPO_VGLUT2 produces a shift in EEG activity consistent with a more activated, wake-like state (Mondino et al., 2021). In the present study, we investigated whether the activation of MLPO_VGLUT2 alters cortical oscillations after sleep deprivation, focusing on delta power and sleep spindles (data are from n = 10 mice) as EEG markers of the NREM (delta power) and REM (theta and sigma power, and spindle density) homeostatic sleep drive (Franken et al., 2001; Borbely et al., 2016; Greene et al., 2017; Bandarabadi et al., 2020; Park et al., 2021). EEG delta power was averaged in 2-hour bins and sleep spindle analysis was conducted on an hour-by-hour basis. The powerful REMs suppression produced by CNO administration precluded any reliable analysis of spectral power during this sleep phase. Statistical analysis was conducted using a mixed-effects model with Geisser-Greenhouse correction, followed by Šídák’s multiple comparisons test. Figures 3A and 3B show, respectively, the mean delta power in the frontal cortex during wakefulness and NREMs, after sleep deprivation. There was a significant effect of treatment (CNO) and time for wakefulness (F(1,10) = 31.44; p = 0.0002 and F(1.597,15.97) = 9.207; p = 0.0035) and NREMs (F(1,10) = 20.05; p = 0.0012 and F(0.9721,9.721) = 6.812; p = 0.0272) delta power; no significant treatment x time interactions were observed (p = 0.9164 and 0.5292). Post hoc multiple comparisons analysis showed that CNO significantly increased EEG delta power during wakefulness (hours 0 - 2, p = 0.0021, hours 2 - 4, p = 0.0006 and hours 4 - 6, p = 0.0019) and NREMs (hours 0 - 2, p = 0.0141, hours 2 - 4, p = 0.0048 and hours 4 - 6, p = 0.0059) following sleep deprivation.

**Figure 3.**
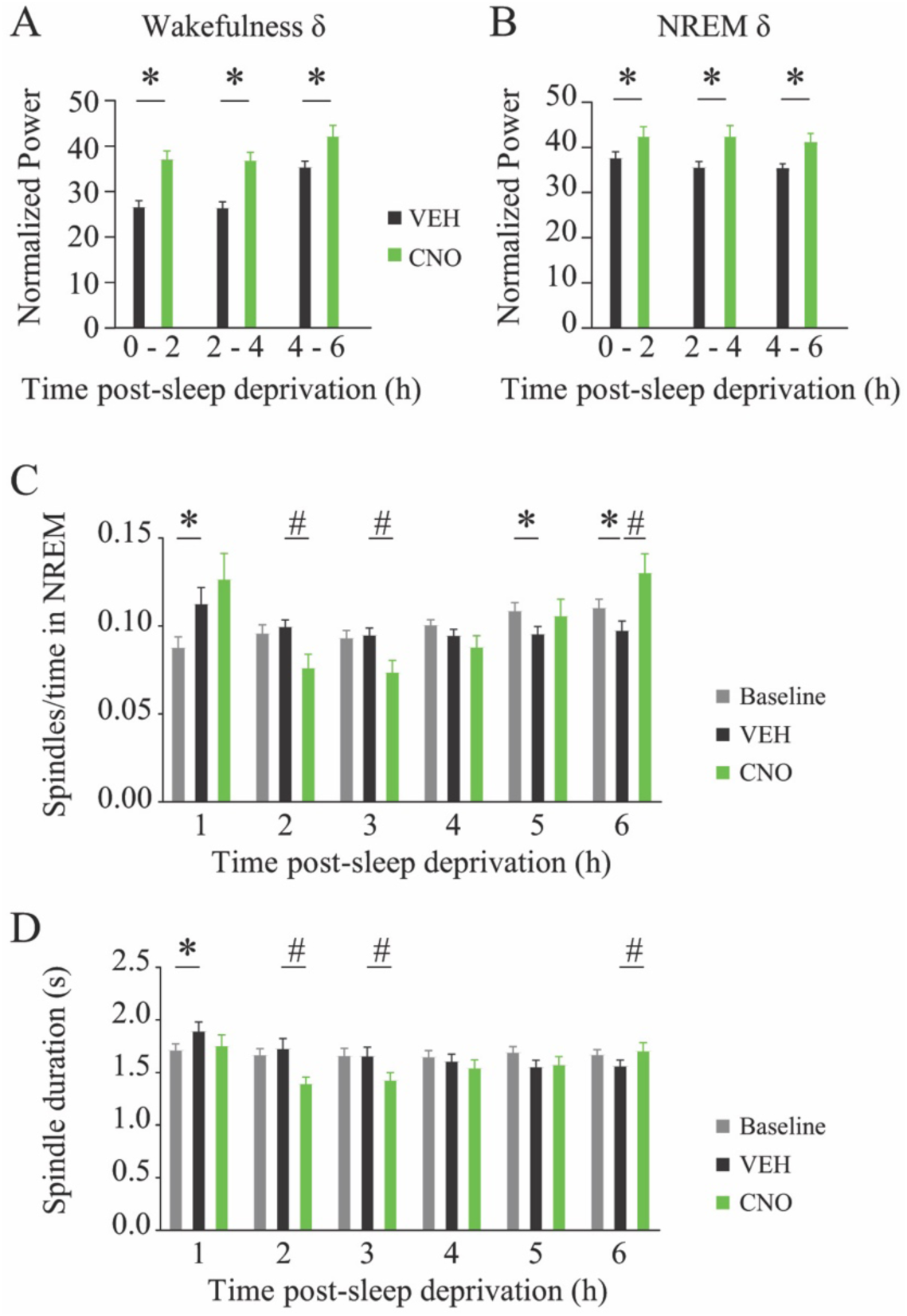
Activation of glutamatergic neurons in the medial-lateral preoptic region altered EEG spectral power after sleep deprivation. **A-B,** Graphs illustrate changes in EEG delta power during wakefulness and NREM sleep states after administration of vehicle (VEH) and clozapine-N-oxide (CNO; 1.0 mg/kg) to sleep deprived mice. **C-D,** Number and duration of sleep spindles and per hour after baseline, and vehicle or CNO injection after sleep deprivation. Data are shown as mean ± SEM. Asterisks in C-D indicate significant differences between VEH and baseline (*, p < 0.05). Hashtags in C-D indicate significant differences between CNO and VEH (^#^, p < 0.05).

Figures 3C and 3D depict the changes in the number and duration of sleep spindles during baseline and after vehicle or CNO injection, quantified during NREMs (no state transitions) after sleep deprivation. Relative to baseline, there was a significant increase in the number and duration of sleep spindles (p = 0.0036 and 0.0082, respectively) during hour 1 of recovery NREMs following sleep deprivation. Furthermore, in comparison with vehicle injection, CNO produced a significant decrease in NREM spindle numbers (p = 0.0150 and 0.0207) and duration (p < 0.0001 and p = 0.0005) during hours 2 and 3 post-sleep deprivation. Both the number of spindles and duration were significantly increased by CNO at hour 6 after sleep deprivation (p = 0.0366 and 0.0474).

### Activation of MLPO Glutamatergic Neurons at Night Suppressed REM Sleep Without Substantially Altering the Time Spent in Wakefulness and NREM Sleep

Previously published data showed that activation of MLPO_VGLUT2 during the day-time increased wakefulness (Vanini et al., 2020; Mondino et al., 2021). In this study, we tested whether the effects of MLPO_VGLUT2 on sleep-wake states vary as a function of the time of the day at which these neurons are stimulated. Thus, mice (n = 6) received an injection of vehicle or CNO before starting sleep recordings at the onset of the dark period (ZT12). Analysis of sleep-wake parameters at night was conducted using a two-way, repeated-measures ANOVA with a Geisser-Greenhouse correction followed by Šídák’s multiple comparisons test. Activation of MLPO_VGLUT2 at night significantly reduced REMs time but did not cause any relevant change (in magnitude and duration) in the time spent in wakefulness and NREMs. There was a significant treatment (CNO) effect on the time spent in REMs and NREMs (F(1,60) = 176.9; p < 0.0001 and F(1,60) = 5.047; p = 0.0284). Post hoc Šídák’s revealed a significant decrease in REMs time between hours 1 and 6 after CNO administration (p = 0.0025, p < 0.0001, p < 0.0001, p = 0.0094, p < 0.0001 and p < 0.0001; Fig. 4A) and an increase in NREMs during hour 2 (p = 0.0002; Fig. 4B). Additionally, there was a significant time effect and treatment x time interaction, but no significant treatment effect on the time spent in wakefulness (F(5,60) = 6.997; p < 0.0001), F(5,60) = 4.852; p = 0.0009 and F(1,60) = 0.1851; p = 0.6685). There was a significant decrease in wakefulness during hour 2 after CNO injection (p = 0.0053; Fig. 4C). Two-way ANOVA showed a significant drug and time effect, as well as drug x time interaction in the number of REMs bouts (F(1,60) = 137.9; p < 0.0001, F(5,60) = 13.22; p < 0.0001 and F(5,60) = 9.561; p < 0.0001).

**Figure 4.**
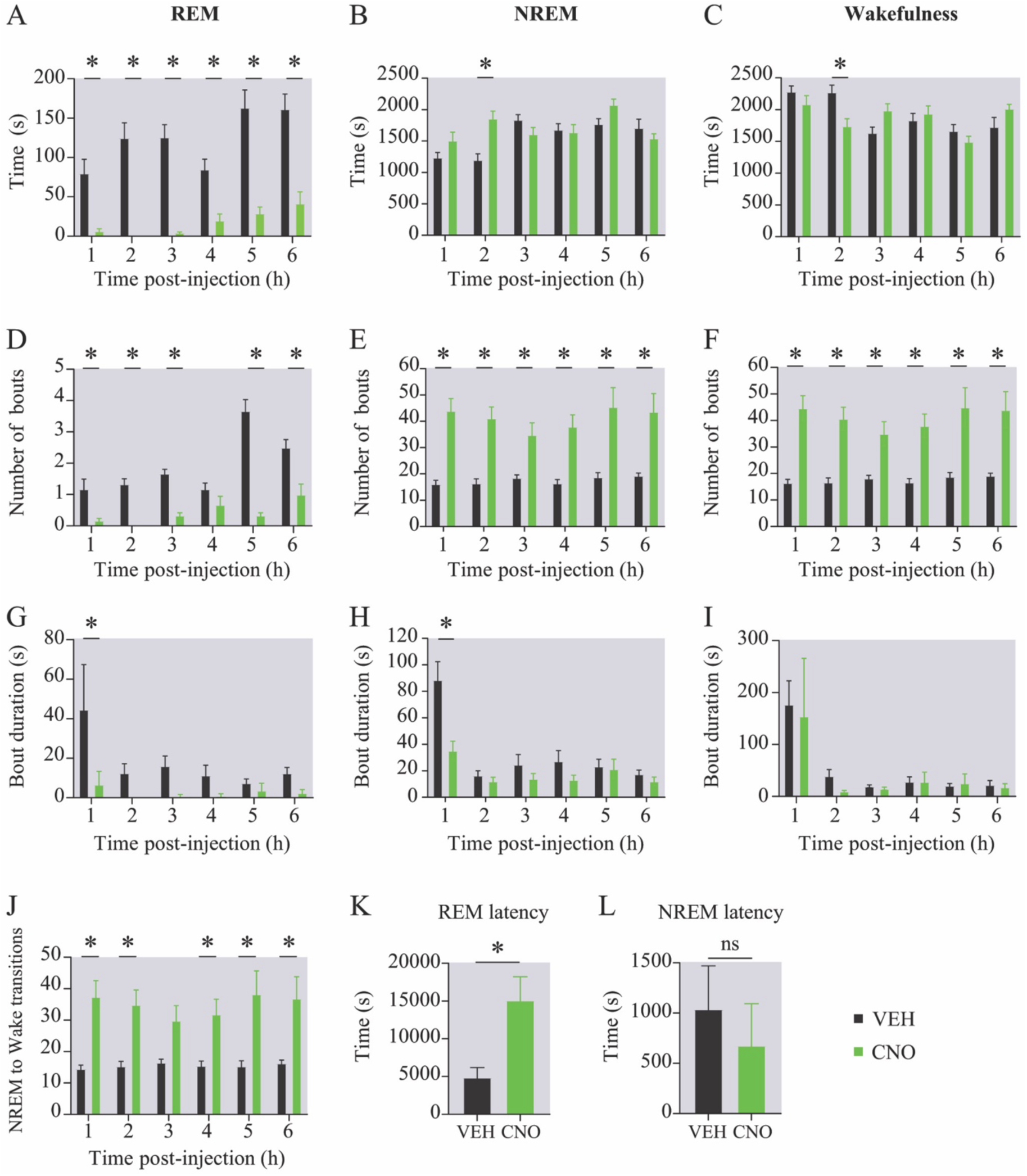
Chemogenetic activation of glutamatergic neurons in the medial-lateral preoptic region during the night period suppressed REM sleep generation. **A-C,** Time spent in REM sleep, NREM sleep, and wakefulness after injection of vehicle (VEH) and clozapine-N-oxide (CNO; 1.0 mg/kg). **D-F,** Number of bouts per state. **G-I,** Mean duration of REM sleep, NREM sleep, and wake bouts. **J,** Number of transitions from NREM sleep to wakefulness. **K-L,** Latency to the first REM and NREM sleep bout after CNO administration. Data are shown as mean ± SEM. The shaded (gray) background in all graphs indicates that the recordings were performed during the night. *, p < 0.05.

Consistent with the decrease in the time spent in REMs, CNO produced a significant decrease in the number of REMs bouts on hour 1, 2, 3, 5 and 6 post-injection (p = 0.0134, p = 0.0005, p = 0.0004, p < 0.0001 and p < 0.0001) (Fig. 4D). Further, ANOVA indicated a significant drug effect on the number of NREMs and wake bouts (F(1,60) = 104.2; p < 0.0001 and F(1,60) = 103.6; p < 0.0001), and no significant time effect or treatment x time interaction (p = 0.6816, p = 0.7452 for NREMs and p = 0.6793, p = 0.7760 for wakefulness). The hour-by-hour analysis showed that CNO significantly increased the number of NREM (p < 0.0001, p = 0.0003, p = 0.0318, p = 0.0020, p < 0.0001 and p = 0.0004) and wake (p < 0.0001, p = 0.0005, p = 0.0255, p = 0.0023, p = 0.0001 and p = 0.0003) bouts between hours 1 and 6 post-injection (Figs. 4E and 4F). Figure 4G-I shows the mean bout duration for REMs, NREMs, and wakefulness after CNO administration. CNO significantly altered bout duration in REMs (F(1,60) = 12.20; p = 0.0009) and NREMs (F(1,60) = 15.98; p = 0.0002). Both REMs (p = 0.0037) and NREMs (p < 0.0001) bout duration significantly decreased during hour 1 post-CNO injection. There was no significant effect on wake episode duration (F(1,60) = 0.2092; p = 0.6491). Relative to vehicle injection, CNO increased the number of transitions from NREM to wakefulness (F(1,60) = 65.16; p < 0.0001) (Fig. 4J). Transitions were significantly increased in hour 1 (p = 0.0014), 2 (p = 0.0087), 4 (p = 0.0409), 5 (p = 0.0014) and 6 (p = 0.0051) post-CNO injection. Consistent with the reduction in REMs bouts after CNO injection, there was a significant increase in the latency to REMs (p = 0.0318; Fig. 4K). However, the latency to NREMs did not change (p = 0.3107; Fig. 4L).

### Activation of MLPO Glutamatergic Neurons at Night Altered EEG Spectral Power During NREM Sleep and Wakefulness

Here we examined the effects of the activation of MLPO_VGLUT2 on cortical oscillations during the night-time period. Frontal cortex EEG spectral power for each frequency band was averaged in 2-hour bins (0-2, 2-4, and 4-6 hours) after vehicle or CNO administration in n = 6 mice. The power spectral analysis focused on wakefulness and NREM states only; the suppression of REMs precluded statistical comparisons of this state. Statistical analysis was conducted using a mixed-effects model with Geisser-Greenhouse correction, followed by Šídák’s multiple comparisons test. Mean changes in EEG power are shown in Figure 5. There was no significant drug effect in wakefulness during hours 0-2 (F(1,30) = 0.5156; p = 0.4783) and 2-4 (F(1,30) = 0.4016; p = 0.5311) after CNO injection, and in NREMs during the first 2 hours post-CNO (F(1,30) = 0.0765; p = 0.7839). Post hoc Šídák’s test indicated that CNO significantly increased delta power in wakefulness during hours 4-6 post-injection (p < 0.0001), and in NREMs during hours 2-4 and 4-6 after CNO administration (p = 0.0246 and p < 0.0001, respectively). In addition, there was a significant decrease in frontal theta power in NREMs during hours 4-6 after CNO injection (p = 0.0036).

**Figure 5.**
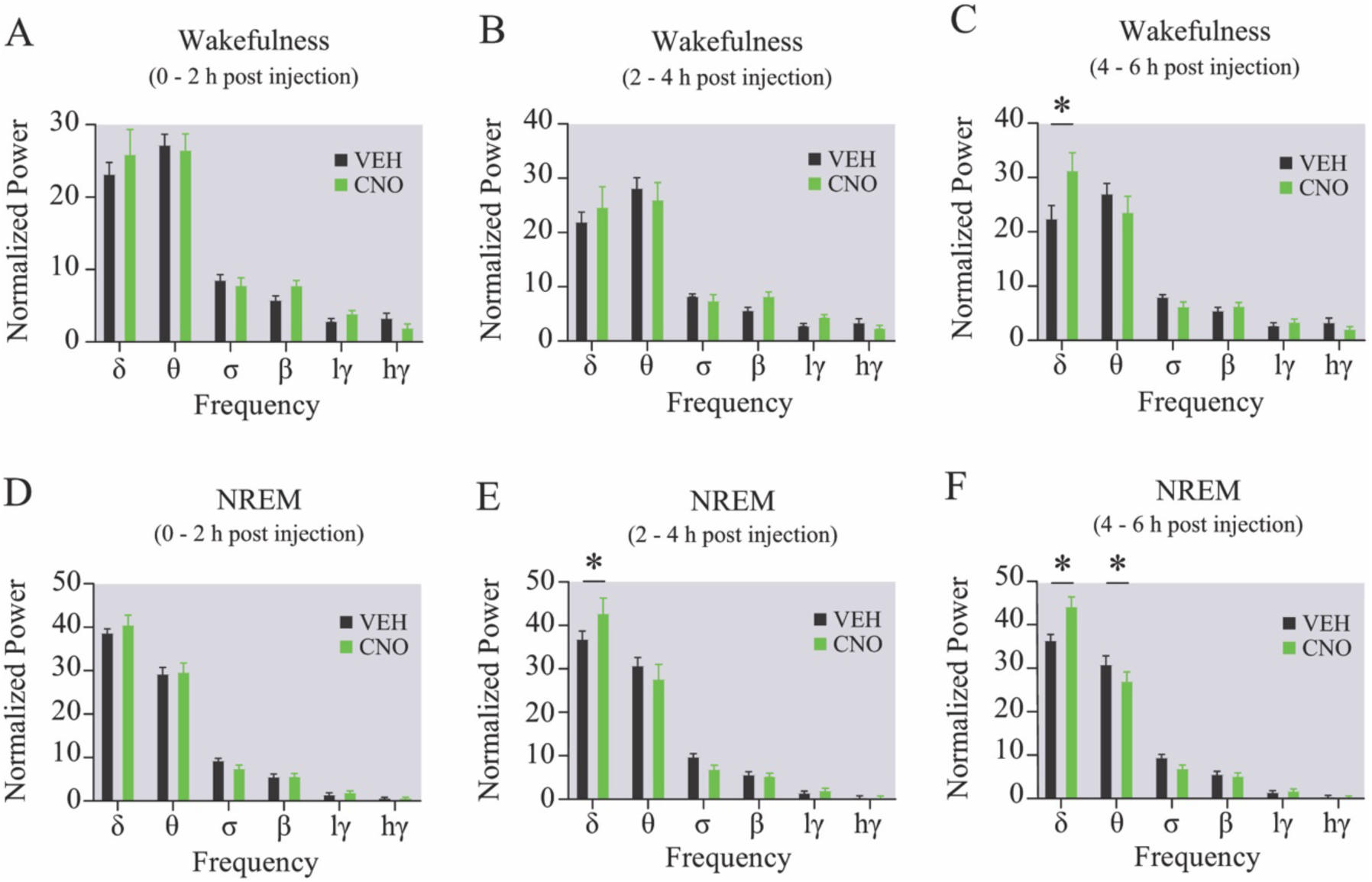
Chemogenetic activation of glutamatergic neurons in the medial-lateral preoptic region during the night period increased EEG delta power in wakefulness and NREM sleep. **A-C,** Graphs show changes in frontal EEG power by frequency band during wakefulness, after injection of vehicle (VEH) and clozapine-N-oxide (CNO; 1.0 mg/kg). **D-F,** Changes in frontal EEG power by frequency band during NREM sleep, after injection of vehicle and CNO. Data are shown as mean ± SEM. All graphs plot mean power analyzed in 2-hour bins post injection. The shaded (gray) background in all graphs indicates that the recordings were performed during the night. *, p < 0.05.

### Inhibition of MLPO Glutamatergic Neurons Promoted REM Sleep Onset But Did Not Alter the Homeostatic Response to Sleep Deprivation

Previous work and this study established that exogenous activation of MLPO_VGLUT2 inhibits REM sleep (Mondino et al., 2021). Here we used selective chemogenetic inhibition (Reitz et al., 2020) in 10 Vglut2-Cre mice to determine whether these neurons are required for REMs control. Schematic AAV injection, representative sleep-wake states, and localization of the area of expression of hM4 designer receptors are shown in Figures 6A-B. The total time spent in REMs, NREMs, and wakefulness is plotted in Figures 6C-E for the light phase, and in Figures 6F-H for the dark phase. Analysis of sleep-wake parameters was conducted using a two-way, repeated-measures ANOVA with a Geisser-Greenhouse correction followed by Šídák’s multiple comparisons test. There was a significant treatment (CNO) and time effect for the time in REM and NREM sleep (F(1,48) = 10.12; p = 0.0026, F(5,48) = 9.351; p < 0.0001 and F(1,48) = 6.464; p = 0.0143, F(5,48) = 3.189; p = 0.0145). Relative to control, inhibition of MLPO_VGLUT2 during the light phase (ZT03 – ZT09) significantly increased the time in REMs between hours 1 and 3 post-CNO administration (p = 0.0116, 0.0243 and 0.0336; Fig. 6C). NREMs time was increased during the first hour (p = 0.0289; Fig. 6D). Two-way ANOVA indicated a significant treatment (CNO) and time effect, as well as treatment x time interaction in the time spent in wakefulness (F(1,48) = 9.154; p = 0.0040, F(5,48) = 4.430; p = 0.0021 and F(5,48) = 2.732; p < 0.0299). Wake time was reduced during hours 1, 2, and 6 post-injection (p = 0.0109, 0.0257, and 0.0373; Fig. 6E). There was no change in the number of sleep-wake bouts. REMs and NREMs bout duration were significantly increased during hour 1 post-CNO injection (p < 0.0001 and 0.0012). CNO administration did not significantly alter the number of NREM to wake transitions. CNO produced a significant decrease in the latency to both REMs (mean latency in s ± SEM = 3225 ± 493 vs 2295 ± 198; p = 0.0147) and NREMs (761 ± 189 vs 280 ± 57; p = 0.0106). During the dark phase (ZT12 – ZT18; the rodent’s active period with lowest sleep drive), chemogenetic inhibition of MLPO_VGLUT2 significantly increased the time spent in REMs during hour 2 only (p = 0.0466; Fig. 6F). CNO administration increased the number of REMs bouts during hour 2 (p = 0.0152), and REMs bout duration during hour 1 (p = 0.0012) post-injection. There was no significant difference in the number of NREM-to-wake transitions, as well as in the latency to the first REMs and NREMs bout during the dark phase.

**Figure 6.**
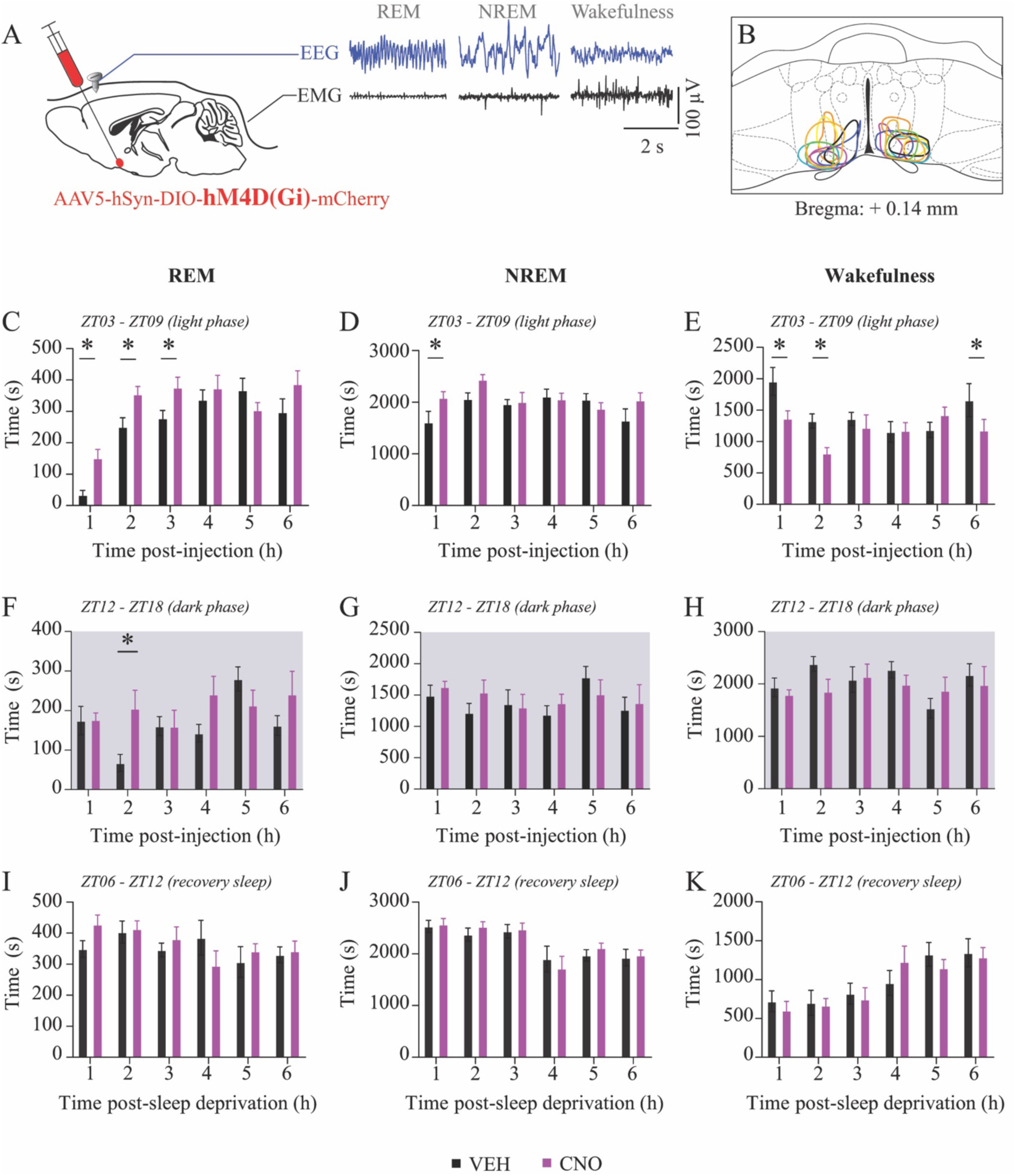
Chemogenetic inhibition of glutamatergic neurons in the medial-lateral preoptic region promoted REM sleep generation during the light phase but did not influence REM sleep homeostasis. **A,** Schematic depiction of the injection of a Cre-dependent adeno-associated virus for expression of the inhibitory designer receptor hM4Di into the medial-lateral preoptic region of Vglut2-Cre mice. Representative recordings of the electroencephalogram (EEG) and electromyogram (EMG) obtained from the same Vglut2-Cre mouse during rapid eye movement (REM) sleep, non-rapid eye movement (NREM) sleep, and wakefulness. **B,** Depiction of the vector injection area, represented on a coronal schematic of the preoptic area modified from a mouse brain atlas (Paxinos and Franklin, 2001). **C-E,** Time spent in REM sleep, NREM sleep, and wakefulness, after administration of vehicle (VEH) and clozapine-N-oxide (CNO; 1.0 mg/kg) to Vglut2-Cre mice during the light phase. Sleep recordings were conducted during the day between ZT03 and ZT09. **F-H,** Graphs summarize the time spent in REM sleep, NREM sleep, and wakefulness, after administration of vehicle (VEH) and clozapine-N-oxide (CNO; 1.0 mg/kg) to Vglut2-Cre mice during the dark phase. Sleep recordings were conducted during the night (indicated by gray background) between ZT12 and ZT18. **I-K,** Time spent in REM sleep, NREM sleep, and wakefulness, after administration of vehicle (VEH) and clozapine-N-oxide (CNO; 1.0 mg/kg) to sleep deprived Vglut2-Cre mice. Sleep deprivation and sleep recordings began, respectively, at ZT0 and ZT06. Data are shown as mean ± SEM. *, p < 0.05.

In this study, we show that chemogenetic stimulation of MLPO_VGLUT2 obliterated the REMs homeostatic response to sleep deprivation (Fig. 2). To investigate whether these neurons play a direct key role in the regulation of sleep homeostasis, Vglut2-Cre mice (n = 10) underwent 6 hours of sleep deprivation and received an injection of vehicle or CNO, followed by 6 hours of sleep-wake recordings. Statistical analysis of changes in sleep-wake parameters after sleep deprivation was conducted using a mixed-effects model with Geisser-Greenhouse correction, followed by Šídák’s multiple comparisons test. Figure 6I-K plots the time spent in REMs, NREMs, and wakefulness after sleep deprivation with and without chemogenetic inhibition of MLPO_VGLUT2. Relative to control, CNO administration did not alter the time spent in REMs, NREMs, and wakefulness after sleep deprivation. Additionally, there was no significant difference after sleep deprivation in the number and duration of bouts of REMs, NREMs, and wakefulness, the number of transitions between NREM and wakefulness, or the latency to REMs and NREMs.

### MLPO Glutamatergic Neurons Send Projections to Arousal-Promoting and REM-Inhibitory Regions

To identify potential neuroanatomic mediators accounting for increased arousal and REMs suppression, we mapped the projections of MLPO_VGLUT2. Mice (n = 5) were injected with AAV- ChR2-mCherry for conditional expression of channelrhodopsin, and brain sections were immunostained for mCherry (Fig. 7). We found dense projections from MLPO_VGLUT2 in the olfactory nucleus, the anterior cingulate cortex, several hypothalamic nuclei (arcuate, periventricular, suprachiasmatic, supramammillary, dorsomedial and ventromedial hypothalamic nuclei), the lateral hypothalamic area, perifornical region, zona incerta, the nucleus accumbens (core and shell), and the thalamus (central medial thalamic and centrolateral thalamic nuclei). MLPO_VGLUT2 also project to the medial septum and caudal portion of the cholinergic basal forebrain (substantia innominata, nucleus of the horizontal limb of the diagonal band of Broca and bed nucleus of the stria terminalis). Furthermore, labeled axons were identified within the periaqueductal gray (dorsal and ventrolateral subdivisions), deep mesencephalic nucleus, the laterodorsal tegmental nucleus, parabrachial nucleus, as well as in several monoaminergic nuclei in the hypothalamus (tuberomammillary nucleus), midbrain (ventral tegmental area, dorsal and median raphe nuclei), pons (locus coeruleus), and medulla (raphe magnus, pallidus and obscurus) regions.

**Figure 7.**
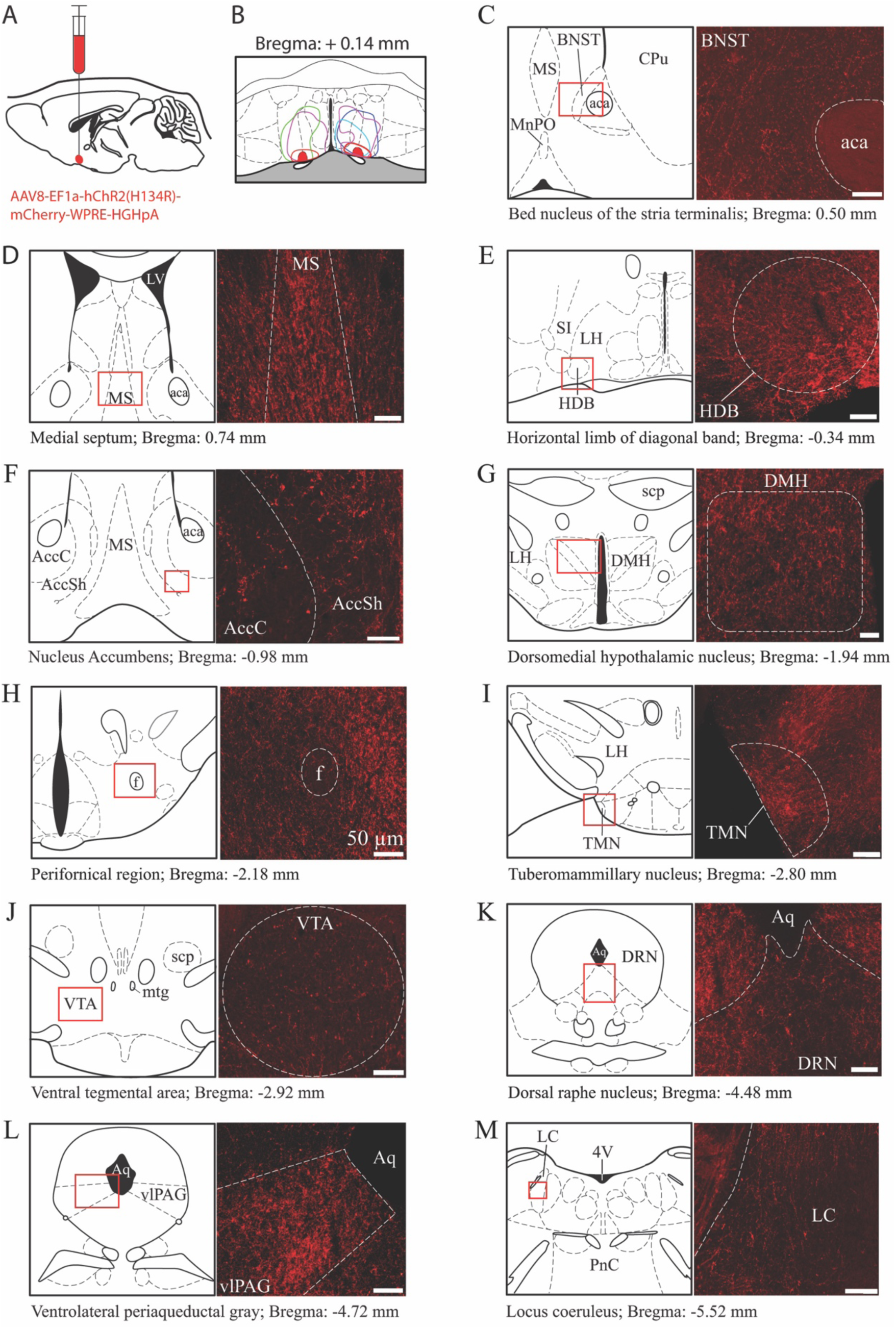
Mapping of efferent projections from glutamatergic neurons in the medial-lateral preoptic region. **A,** Sagittal view of the mouse brain schematizing the injection of the adeno-associated virus AAV-ChR2-mCherry for conditional expression of channelrhodopsin in medial-lateral preoptic glutamatergic neurons and their axonal projections. **B,** Color-coded depiction of the area of channelrhodopsin expression within the medial-lateral preoptic region (vector injection site) is represented on a coronal schematic of the preoptic area modified from a mouse brain atlas (Paxinos and Franklin, 2001). **C-M,** mCherry-positive (red) axonal projection targets of glutamatergic neurons in the medial-lateral preoptic region are shown on selected coronal sections containing several basal forebrain, hypothalamic, and pontine regions. Photographs are from the area indicated by the red squares on each schematic coronal view of brain and brain stem sections. Scale bars, 50 µm. Abbreviations: aca, anterior commissure; AccC, nucleus accumbens core; AccSh, nucleus accumbens shell; Aq, aqueduct; BNST, bed nucleus of the stria terminalis, CPu, striatum; DMH, dorsomedial hypothalamic nucleus; DRN, dorsal raphe nucleus; f, fornix; HDB, nucleus of the horizontal limb of the diagonal band, LC, locus coeruleus; LH, lateral hypothalamus; LV, lateral ventricle; MnPO, median preoptic nucleus; MS, medial septum; mtg, mammillotegmental tract; PnC, pontine reticular nucleus, caudal part; SI, substantia innominate; scp, superior cerebellar peduncle; TMN, tuberomammillary nucleus; vlPAG, ventrolateral periaqueductal gray; VTA, ventral tegmental area; 4V, fourth ventricle.

## DISCUSSION

This study revealed that MLPO_VGLUT2 play a critical role in regulating spontaneous REMs as well as the REMs homeostatic response to sleep deprivation. In fiber photometry recordings, we found that MLPO_VGLUT2 are maximally active during REMs and brief arousals from sleep, and exhibit minimally increased activity during wake bouts. These results partly agree with recent work showing that preoptic glutamatergic neurons are wake-active and promote wakefulness (Mondino et al., 2021; Smith et al., 2024). Notably, our finding that these neurons are REMs-active was not reported in previous photometry recording studies of preoptic glutamatergic neurons. This discrepancy could be explained by the different locations of our recording sites, which were more rostral relative to the stereotaxic target sites reported in *Smith et al., 2024* (+0.14 versus +0.40mm from Bregma). Additionally, we cannot rule out that our data are comprised of two separate neuronal populations. However, our results are consistent with several reports of wake/REM active neurons distributed across numerous forebrain (Hassani et al., 2009), hypothalamic (including preoptic nuclei) (Sakai, 2011), and brainstem sites (Steriade et al., 1990; Lee et al., 2001; Boucetta et al., 2014; Cisse et al., 2018; Sakai, 2018). Future studies are crucial to elucidate (1) the precise circuitry by which MLPO_VGLUT2 influence REMs and wakefulness, (2) whether there are two functionally distinct groups of MLPO_VGLUT2, and (3) whether the REMs- promoting glutamatergic neurons in the sublaterodorsal nucleus (Boissard et al., 2002; Clement et al., 2011) are indirectly inhibited by the REM-active group (discharges to end REMs) or the wake-active group (discharges to block REMs onset).

Next, we show that chemogenetic stimulation of MLPO_VGLUT2 impaired the homeostatic response to sleep deprivation. The disruption of sleep homeostasis was characterized by a slight, transient increase in wakefulness that delayed (by 1 hour) the homeostatic rebound of NREMs. However, the most salient and relevant effect was a near-complete suppression of REMs during the 6-hour sleep recovery phase. Additionally, the activation of MLPO_VGLUT2 increased EEG delta power during NREMs and wake states after sleep deprivation. The increase in delta power, relative to that observed in mice that underwent sleep deprivation without neuronal stimulation, suggests that MLPO_VGLUT2 further increased sleep propensity and interfered with cortical homeostatic processes. These results are consistent with previously published data supporting the conclusion that activation of MLPO_VGLUT2 promotes wakefulness and reduces NREMs (Mondino et al., 2021). However, the present work is novel because no studies have investigated how these wake-promoting preoptic neurons influence sleep homeostasis. Another novel finding of our study was that MLPO_VGLUT2 send efferent projections to central regions that promote wakefulness, suggesting that these neurons can indirectly influence cortical and behavioral arousal via excitatory inputs to several subcortical arousal-promoting systems. Further, this study extends previous findings to suggest a more specialized function of these neurons controlling REMs generation, even when the homeostatic sleep drive is high.

Another important effect produced by the activation of MLPO_VGLUT2 was a sustained increase in the number of NREMs bouts and NREM-to-wake transitions following sleep deprivation. Together with the increase in the number of wake bouts, this is indicative of sleep fragmentation, which we have reported previously after activation of these preoptic cells in non-sleep deprived mice (Mondino et al., 2021). The NREM state fragmentation is robust, despite the expected additive effect of sleep deprivation during the early light period when the sleep drive is normally highest. We propose that NREMs fragmentation is one potential underlying cause for the increase in NREM and wake delta power, observed after sleep deprivation with the activation of MLPO_VGLUT2. Sleep fragmentation precludes NREMs consolidation and would thus produce an incomplete recovery from the sleep debt accrued during sleep deprivation, which is reflected by the increase in delta power during subsequent wakefulness and NREMs. These results are biologically and translationally relevant because sleep fragmentation is a prominent feature of the abnormal sleep patterns observed in sleep apnea (Guilleminault et al., 1976; Kimoff, 1996), neuropsychiatric disorders (Lim et al., 2014), and aging (Lim et al., 2014; Li et al., 2018).

Importantly, the number of galaninergic neurons in the intermediate nucleus – the human homologue of the mouse ventrolateral preoptic nucleus – has been shown to correlate with sleep fragmentation and sleep loss in aging and Alzheimer’s disease (Lim et al., 2014). Furthermore, a recent study by Li and colleagues (Li et al., 2022) demonstrated that hypocretin/orexin neurons become hyperexcitable and contribute to fragmented sleep in aging mice. Thus, future studies of the interactions between MLPO_VGLUT2, ventrolateral preoptic galaninergic, and hypocretin/orexin neurons will be important for advancing our understanding of the mechanisms underlying disrupted sleep-wake cycling during aging. Our results also have translational potential because, in addition to the sleep fragmentation, activation of MLPO_VGLUT2 caused a reduction in the number and duration of sleep spindles after sleep deprivation. Both sleep fragmentation and abnormal spindle oscillations are detrimental to cognitive function (Mednick et al., 2013; Astill et al., 2014; Maski et al., 2017; Swift et al., 2018; Varela and Wilson, 2020; Dickey et al., 2021; Petzka et al., 2022). Future work will be required to assess the impact of MLPO_VGLUT2 on different cognitive domains, especially attention and memory consolidation.

To further characterize the role of MLPO_VGLUT2 in the regulation of sleep-wake states, a set of stimulation experiments conducted at night revealed no substantial changes in NREM and wake states. These results were predictable because the activation of these neurons during the light period generates a modest increase in wakefulness, and so their wake-promoting effects during the rodent’s active phase – when the sleep drive is lowest – are expected to be confounded by circadian factors. Sleep fragmentation and REMs suppression were, again, the main significant changes and appear to be the most consistent and reproducible effects across studies. Activation of MLPO_VGLUT2 at night also increased EEG delta power during wakefulness and NREMs. The increase in delta power that occurred towards the second half of the sleep study suggests that, possibly through sleep fragmentation, MLPO_VGLUT2 can enhance the NREMs debt accrued during the active night period.

Chemogenetic inactivation of MLPO_VGLUT2 produced a sustained increase (3 hours) in REMs. Relative to the modest and short-lasting effect observed in NREMs, the substantial increase in REMs supports the notion that MLPO_VGLUT2 play a specific role in the regulation of REMs. Interestingly, chemogenetic inactivation did not alter sleep-wake time during the dark period, indicating that the activity of MLPO_VGLUT2 is under a strong circadian influence. Afferent projections to the preoptic area have been extensively studied, but specific synaptic inputs to MLPO_VGLUT2 remain to be identified. The lack of substantial changes in sleep-wake architecture observed after sleep deprivation suggests that these MLPO_VGLUT2 play a key role in normal physiologic REMs control but may not be critical for REMs homeostasis.

Several lines of evidence suggest that subsets of sleep-active preoptic neurons may promote REMs (Lu et al., 2002; Suntsova et al., 2002; Gvilia et al., 2006; Takahashi et al., 2009; Sakai, 2011; Alam et al., 2014). In contrast, here and in previous studies (Vanini et al., 2020; Mondino et al., 2021) we show that MLPO_VGLUT2 exert a potent inhibitory control on REMs generation. Although there are several potential mechanisms underlying this effect (see detailed discussion in Mondino et al., 2021), here we demonstrate that MLPO_VGLUT2 project to several regions of the hypothalamus and brainstem including the perifornical region of the hypothalamus (Lu et al., 2007; Watson et al., 2008; Brevig et al., 2010; Kummangal et al., 2013; Torterolo et al., 2013), the ventrolateral periaqueductal gray (Sastre et al., 1996; Vanini et al., 2007; Sapin et al., 2009; Weber et al., 2018), as well as several monoaminergic nuclei that directly inhibit REMs generation (Sakai and Crochet, 2000; Pal and Mallick, 2006; Khanday et al., 2016; Saito et al., 2018; Osorio-Forero et al., 2021; Fenik and Rukhadze, 2022).

Collectively, our study revealed that MLPO_VGLUT2 are mostly active during spontaneous REMs and wakefulness. Chemogenetic experiments suggest that MLPO_VGLUT2 are necessary and sufficient for the control of spontaneous REMs generation, but only the activation of these neurons appears to impair REMs homeostasis after sleep deprivation. Importantly, expression of the calcium sensor and designer receptors spared adjacent basal forebrain sites containing glutamatergic neurons that promote wakefulness (Xu et al., 2015). Further, this study identified downstream projections to several arousal systems through which MLPO_VGLUT2 presumably promote wakefulness and suppress REMs. Future studies are essential to establish the individual contribution of these pathways to the control of sleep-wake cycles. The present work is limited by the absence of photometry data during sleep deprivation/recovery, the study of sex-specific responses, and the possibility that – due to the rich cellular diversity of the preoptic region (Moffitt et al., 2018) – two different functional groups of MLPO_VGLUT2 were simultaneously targeted by chemogenetic manipulations.

Last, our results also suggest that state-specific activation of MLPO_VGLUT2 may be critical for creating a barrier between REMs and wakefulness, which – when considered from the evolutionary perspective – would be critical after arousal from sleep to ensure fully connected consciousness directed to environmental events and the prevention of REMs intrusion, which would limit cognitive and motoric function in response to threat.

## Acknowledgments

This work was funded by the National Institutes of Health (R01 GM124248 and R01 AG078134 to G.V. and G.A.M.), Bethesda, MD, USA, and the Department of Anesthesiology, University of Michigan Medical School, Ann Arbor. We thank Dr. Chris Andrews, Ph.D., (Consulting for Statistics, Computing & Analytics Research, University of Michigan, Ann Arbor, Michigan) for help with statistical analysis, and Dr. James M. McNally from the Department of Psychiatry, VA Boston Healthcare System and Harvard Medical School for providing his automated sleep spindle detection algorithm.

## Notes

### Competing Interest Statement

The authors have declared no competing interest.

